# Different brain systems support the aversive and appetitive sides of human pain-avoidance learning

**DOI:** 10.1101/2021.10.18.464769

**Authors:** Marieke Jepma, Mathieu Roy, Kiran Ramlakhan, Monique van Velzen, Albert Dahan

## Abstract

Both unexpected pain and unexpected pain absence can drive avoidance learning, but whether they do so via shared or separate neural and neurochemical systems is largely unknown. To address this issue, we combined an instrumental pain-avoidance learning task with computational modeling, functional magnetic resonance imaging (fMRI) and pharmacological manipulations of the dopaminergic (100 mg levodopa) and opioidergic (50 mg naltrexone) systems (N=83). Computational modeling provided evidence that untreated participants learned more from received than avoided pain. Our dopamine and opioid manipulations negated this learning asymmetry by selectively increasing learning rates for avoided pain. Furthermore, our fMRI analyses revealed that pain prediction errors were encoded in subcortical and limbic brain regions, whereas no-pain prediction errors were encoded in frontal and parietal cortical regions. However, we found no effects of our pharmacological manipulations on the neural encoding of prediction errors. Together, our results suggest that human pain-avoidance learning is supported by separate threat- and safety-learning systems, and that dopamine and endogenous opioids specifically regulate learning from successfully avoided pain.

## Introduction

Learning to avoid actions that cause damage to our body is critical for health and survival. The experience of pain is an important teaching signal in this learning process, such as when a child learns to avoid touching a hot stove, or when a patient who underwent knee surgery learns to avoid bending his or her knee. However, the *absence* of otherwise expected pain can be an equally important teaching signal. When, for example, some weeks after surgery a patient realizes that bending his or her knee is *not* painful anymore, this suggests that particular movements are safe again and no longer needs to be avoided. Adaptive behavior in situations associated with pain thus requires an optimal balance between threat and safety learning when confronted with, respectively, the unexpected presence and absence of pain.

Previous studies have made considerable progress in our understanding of the neural basis of passive cue-pain-association learning (Ploghaus et al. 2000, Seymour et al. 2004, Seymour et al. 2005) and—more recently—active pain-avoidance and -relief learning (Roy et al. 2014, Eldar et al. 2016, Zhang et al. 2018) in humans. A key aspect of these studies was the application of reinforcement-learning models to the analysis of neuroimaging data. According to reinforcement-learning theory, learning is driven by *prediction errors*, which signal the difference between the actual and expected outcomes of an action (Sutton and Barto 1998). Thus, actions that result in the unexpected presence versus absence of pain yield oppositely signed prediction errors which, respectively, increase and decrease the aversive value associated with that action. However, whether these opponent teaching signals drive learning via one underlying brain system, or via separate ones, is still largely unknown.

One possibility is that prediction errors elicited by the unexpected presence and absence of pain are encoded as opposite activity patterns in the same brain regions (i.e., one learning system). If this is the case, we may also expect that—at the neurochemical level— learning from these two outcomes is supported by the same neuromodulator(s). Furthermore, these two outcomes may then be equally effective in driving learning, such that they are associated with the same learning rate. Most previous studies, including our own, have assumed that this is the case. For example, in a previous functional magnetic resonance imaging (fMRI) study, we identified brain activity encoding *general* aversive prediction errors, signaling the degree to which *both* received- and avoided-pain outcomes are relatively worse (or less good) than expected (Roy et al. 2014). Another possibility, however, is that learning from received and successfully avoided pain are subserved by two separate brain systems. In this case, learning from these two outcomes may also be supported by different neuromodulatory systems, and associated with different learning rates. Although the notion of two systems is broadly consistent with theoretical accounts of avoidance learning, such as two-factor theory (Mowrer 1951), there is not much empirical evidence for this idea, especially not in humans.

Regarding the role of neuromodulators, there is a wealth of evidence that reward prediction errors, which are thought to drive reward-pursuit behaviors, are signaled by the phasic activity of midbrain dopamine neurons (Schultz et al. 1997) that project to the ventral striatum (O’Doherty et al. 2003, Rutledge et al. 2010). Whether the dopamine system also has a role in aversive learning is controversial, but several hypotheses have been proposed, mostly based on animal studies (Palminteri and Pessiglione 2017). According to one prominent account, the same dopaminergic prediction-error response that reinforces actions associated with reward also reinforces actions associated with the omission of aversive outcomes (Mowrer 1956, Dinsmoor 2001, Moutoussis et al. 2008). This account thus predicts a role for dopamine in learning from successfully avoided pain. However, animal studies have provided mixed evidence for this idea (Josselyn et al. 2005, Oleson et al. 2012, Wietzikoski et al. 2012, Dombrowski et al. 2013, Fernando et al. 2013, Salinas-Hernandez et al. 2018, Wenzel et al. 2018, Stelly et al. 2019) and evidence in humans is scarce (Raczka et al. 2011).

The endogenous opioid system is another interesting candidate neuromodulatory system for pain-avoidance learning. Pavlovian fear conditioning studies in animals have provided evidence for a causal role of opioidergic activity—specifically via μ-opioid receptors in the periaqueductal gray (PAG)—in both fear conditioning (McNally and Cole 2006, Cole and McNally 2007, Cole and McNally 2009, McNally et al. 2011) and fear-extinction learning (McNally and Westbrook 2003, McNally et al. 2004, McNally et al. 2005, Kim and Richardson 2009, Parsons et al. 2010). These findings suggest that the endogenous μ-opioid system may mediate learning from received and/or avoided pain, although this remains to be studied in instrumental learning tasks and in humans.

In the present study, we aimed to dissociate learning from received and avoided pain in terms of behavior (learning rates), neural encoding of prediction errors, and the roles of dopamine and endogenous opioids. To this end, we combined an instrumental pain-avoidance learning task with computational modeling, fMRI, and pharmacological manipulations of the dopamine and opioid systems, in a randomized, double-blind, between-subject design. Specifically, participants completed the learning task under one of three drug conditions: 100 mg levodopa (a dopamine precursor), 50 mg naltrexone (an opioid antagonist, with highest affinity for the μ-opioid receptor), or placebo. PET studies in humans suggest that levodopa increases phasic dopamine bursts (Floel et al. 2008), but not tonic dopamine activity (Black et al. 2015). Thus, if dopaminergic prediction-error responses support learning from successfully avoided pain, we expect levodopa to enhance learning rates and neural prediction error signaling when pain is avoided. Naltrexone blocks the majority of μ-opioid receptors in the brain (Lee et al. 1988, Preston and Bigelow 1993, Schuh et al. 1999, Weerts et al. 2013). Thus, if μ-opioid receptor activity supports learning from received and/or avoided pain outcomes, we expect naltrexone to reduce learning rates and neural prediction error signaling for the corresponding outcome(s).

## Results

Eighty-three healthy human participants completed a pain-avoidance learning task during fMRI, under one of three treatment conditions (levodopa, naltrexone, or placebo). On each of 144 trials of the pain-avoidance learning task, participants chose between two options (Figure 1A), each probabilistically associated with the delivery of painful heat (49 or 50°C, 1.9 s duration) to their left lower leg. Pain probabilities for each option were governed by two independently varying random walks (Figure 1B). Thus, participants had to track the changing reinforcement values, and kept experiencing prediction errors throughout the task.

**Figure 1.**
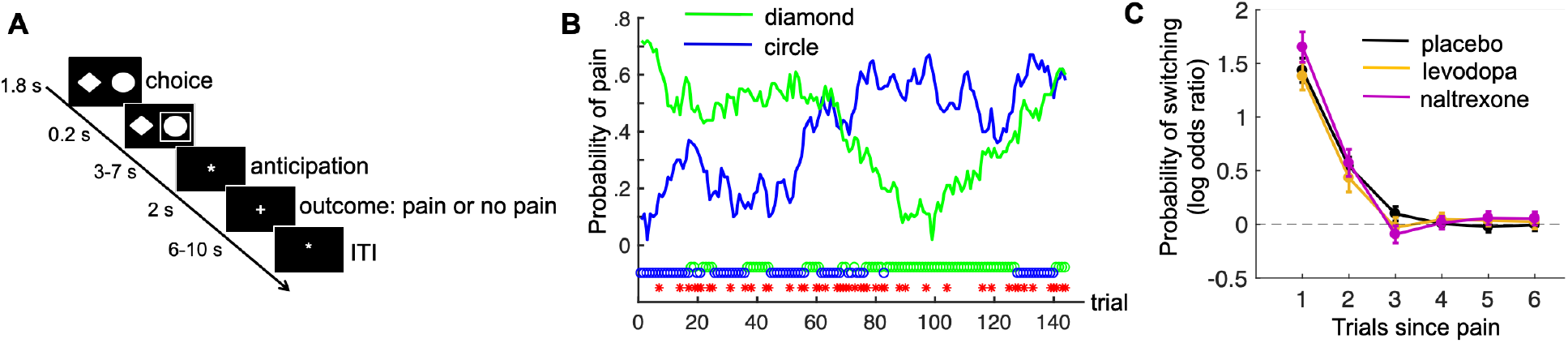
Pain-avoidance learning task. **A.** Outline of one trial. **B**. Example of pain probabilities and choice data for one participant. The green and blue lines show the trial-specific probabilities of receiving a painful heat stimulus when choosing each option. The green and blue circles below the graph indicate the participant’s choices, and the red stars indicate trials on which pain was delivered. **C**. Probability of switching per treatment group, as a function of pain 1-6 trials back. Error bars are standard errors.

There were no treatment effects on subjective state (alertness, calmness or contentment; Figure 1—figure supplement 1) or self-reported heat pain during a separate pain-rating task immediately preceding the avoidance-learning task (Figure 1—figure supplement 2).

Nine participants were excluded from the fMRI (but not the behavioral) analyses because of excessive head movement. Thus, the behavioral analyses included 83 participants (26-29 per treatment group), and the fMRI analyses included 74 participants (24-26 per treatment group).

### No drug effects on model-independent measures of task performance

On average, participants received pain on 57.1 of the 144 trials (SD = 8.6). As expected, participants switched to the other choice option more frequently after receiving pain (46.6% of those trials, SD = 21.2) than after avoiding pain (5.4% of those trials, SD = 6.9; *t*(82)= 18.6, *p <* .001). The effect of previous pain outcomes on switching also decayed exponentially over time, in all treatment groups (Figure 1C; *p <* 0.001 for 1 trial back, *p* <.002 for 2 trials back, and p > .047 for 3-6 trials back, in all groups).

The three treatment groups did not differ in the number of received pain stimuli (*F*(2,80) = 0.56, *p =* 0.57), frequency of switching following pain outcomes (*F*(2,80) = 0.03, *p =* 0.97), or frequency of switching following no-pain outcomes (*F*(2,80) = 1.18, *p =* 0.31). Thus, our pharmacological manipulations did not affect basic measures of task performance.

### Computational modeling

To formalize and quantify the latent learning and decision processes thought to underlie participants’ choice behavior, we applied two candidate reinforcement-learning models to the choice data, using a hierarchical Bayesian approach. Group-level parameters were estimated separately for each treatment group. Both models update the expected pain probability for the chosen option on each trial, in proportion to the prediction error (Rescorla and Wagner 1972). The two models differ in that Model 1 uses a single learning rate, *α*, for all outcomes, whereas Model 2 uses separate learning rates for received and avoided pain: *α*_*pain*_ and *α*_*no-pain*_, respectively (see Methods for model equations and parameter-estimation details). If Model 2 is better able to explain the choice data than Model 1, this could be taken as initial support for the idea that learning from received and avoided pain is subserved by different learning systems.

Both learning models were combined with a softmax decision function that translates expected pain probabilities into choice probabilities. Inverse-temperature parameter *β* controls the degree of choice randomness, such that the likelihood that the model chooses the option with the lowest expected pain probability increases as *β* increases.

#### Model comparison: evidence for asymmetric learning from received and avoided pain

In all treatment groups, the model with separate learning rates for received and avoided pain (Model 2) outperformed the model with a single learning rate (Model 1), providing initial evidence for the presence of two learning systems. The Watanabe-Akaike information criterion (WAIC) for Model 1 vs. Model 2 was 3109 vs. 2959, 2958 vs. 2941, and 3164 vs. 3106 for the placebo, levodopa and naltrexone groups, respectively.

#### Parameter estimates: levodopa and naltrexone increase learning rates for avoided pain

We next examined the parameter estimates of the best fitting model. We focus on the hyperparameters governing the means of the group-level distributions, which we denote with overbars (e.g., 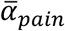 refers to the group-level mean of *α*_*pain*_). Figure 2A shows the posterior distribution of these group-level mean parameters, per treatment group. The corresponding 95% highest density intervals (HDIs) are reported in Figure 2—figure supplement 1. The 95% HDIs of each individual participant’s learning rate posteriors are shown in Figure 2— figure supplement 2.

**Figure 2.**
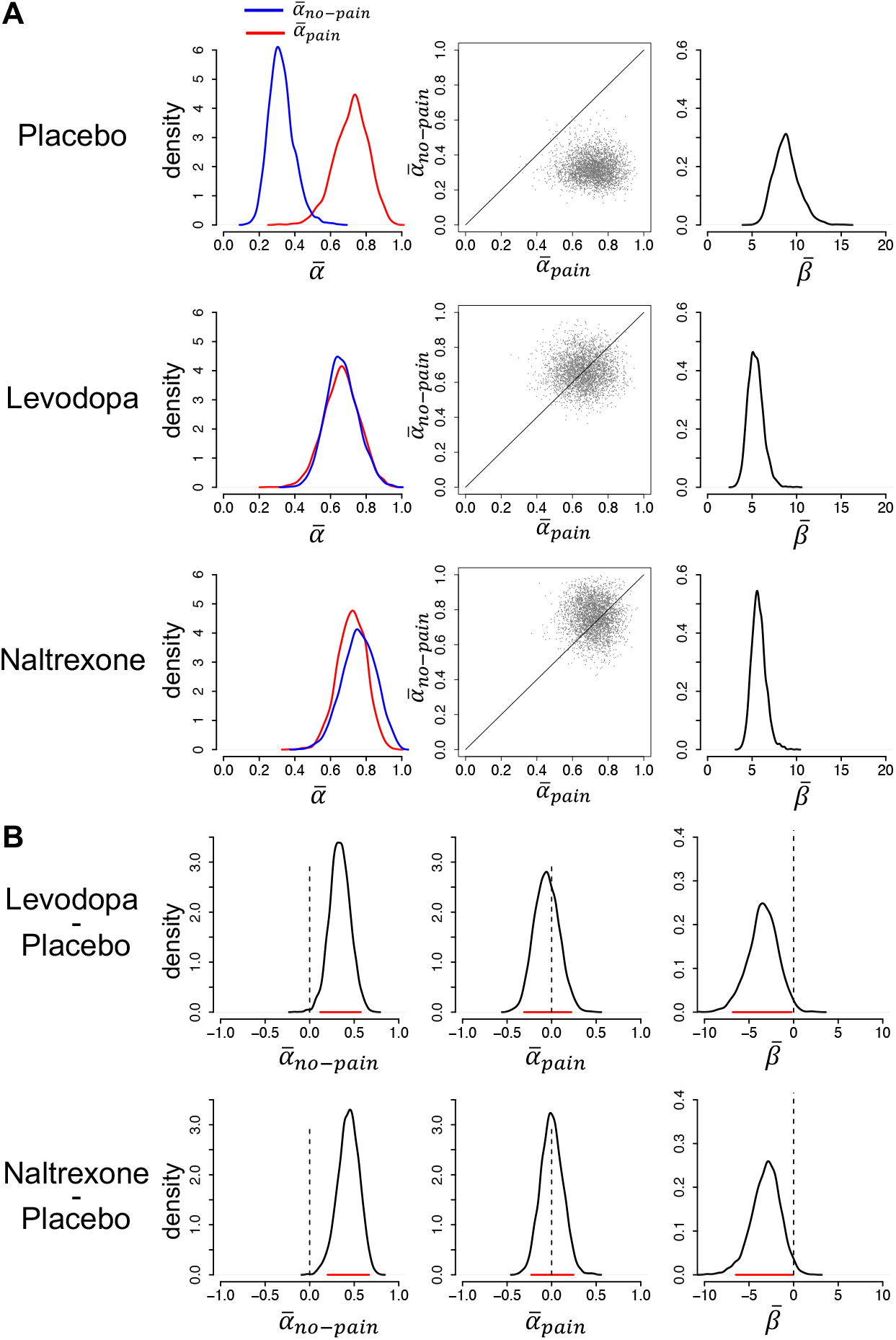
Model parameters. **A**. Posterior distributions of the parameters’ group-level means for each group (left and right panels). Parameters *α*_*no-pain*_ and *α*_*pain*_ are learning rates for avoided and received pain outcomes, respectively; parameter *β* is the inverse-temperature parameter. The middle panels are joint density plots of 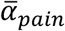 and 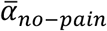 (dots are samples from the MCMC), showing that 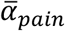 is reliably greater than 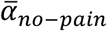 in the placebo group only. **B**. The difference between the posterior distributions for each drug group vs. the placebo group, showing that 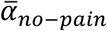 is greater and 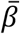 is smaller in both drug groups compared to the placebo group. Red lines indicate 95% HDIs.

In the placebo group, the posterior distribution of 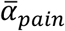 (median = 0.72) was considerably higher than the posterior distribution of 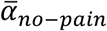 (median = 0.32; 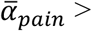 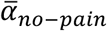 for 99.6% of the MCMC samples), indicative of stronger expectation updating when pain was received than avoided. In contrast, in both drug groups, the posterior distributions of 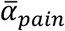 and 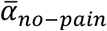 were highly similar, due to a specific increase in 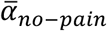 relative to the placebo group. In the levodopa group, the posterior medians of 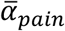 and 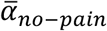 were, respectively, 0.66 and 0.66 (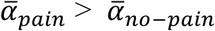 for 50% of the MCMC samples). In the naltrexone group, the posterior medians of 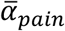 and 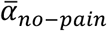 were, respectively, 0.72 and 0.76 (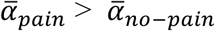 for 38% of the MCMC samples). Note that the best-fitting model for both drug groups contained separate learning rates for received and avoided pain. Combined with the finding that the group-level mean learning-rate parameters for these two outcomes were highly similar, this suggests that some participants in each drug group learned more from received than avoided pain while others showed the opposite bias, but that there was no systematic learning asymmetry (at the individual level, *α*_*pain*_ was higher than *α*_*no-pain*_ for 50% of the levodopa and 41% of the naltrexone participants).

Thus, at the group level, both levodopa and naltrexone, as compared to placebo, increased learning rates for avoided pain, while not affecting learning from received pain (Figure 2A, left and middle panels). To test the significance of these group differences, we computed the difference between the posterior distributions of the group-level mean parameters for each drug group vs. the placebo group (Figure 2B). For 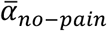, 99.7% of the difference distribution for levodopa vs. placebo, and 99.9% of the difference distribution for naltrexone vs. placebo, lay above 0. In contrast, 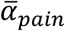 did not differ between the drug and placebo groups: 34% and 49% of the difference distributions for levodopa vs. placebo and naltrexone vs. placebo, respectively, lay above 0. Thus, both drugs selectively increased learning rates for avoided pain.

The posterior distribution of inverse-temperature parameter 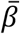 was higher for the placebo group (median = 8.8) than the levodopa and naltrexone group (median = 5.3 and 5.7, respectively), as well. Specifically, 98.8% of the 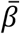 difference distribution for levodopa vs. placebo, and 98.1% for naltrexone vs. placebo, lay below 0, indicating that 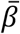 was reliably lower for both drug groups compared to the placebo group. This suggests that participants in the two drug groups, as compared to the placebo group, were less prone to choose the option with the lowest expected pain probability (i.e., more stochastic choice behavior).

Together, the parameter estimates suggest that (i) untreated (placebo group) participants updated their expectations more rapidly following received than avoided pain, (ii) levodopa and naltrexone negated this learning asymmetry by selectively increasing learning rates for avoided pain, and (iii) levodopa and naltrexone also increased choice stochasticity, possibly reflecting a more exploratory or risky choice strategy.

#### Replication of learning rate asymmetry in an independent group of untreated participants

To examine the robustness of our finding of asymmetric learning rates in the placebo group, we applied our hierarchical Bayesian modeling approach to the choice data of 23 untreated participants from our previous pain-avoidance learning fMRI study (Roy et al. 2014), and tested whether the higher learning rate for received than avoided pain found in our placebo group was replicated in this previous dataset (Figure 2—figure supplement 3). In this previous dataset, 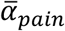 (median = 0.62) was indeed higher than 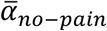 (median = 0.44), resembling the placebo-group results from our current study, although the learning-rate asymmetry was somewhat smaller in the previous dataset (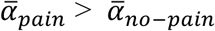 for 90% of the MCMC samples). The posterior median of 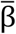 in the previous dataset was 7.1 (95% HDI = 4.7-9.9). The finding of a higher learning rate for pain than no-pain outcomes in this independent dataset corroborates the idea that people normally (in the absence of a pharmacological manipulation) update their expectations more rapidly following received than avoided pain.

#### Parameter recovery

To validate the conclusion that our pharmacological manipulations affected both 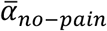 and 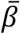, we simulated two sets of choice data, using the parameter values found in the placebo and drug groups, and performed parameter-recovery analyses (Appendix 1). In sum, we found that our modeling procedure can distinguish the two patterns of parameter values found in the placebo and drug groups, even though they produce similar model-independent performance measures (number of received pain stimuli, and frequency of switching following pain and no-pain outcomes). Thus, the observed drug effects on both 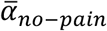 and 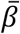 are unlikely to merely reflect a tradeoff between these parameters. Instead, the parameter-recovery results suggest that levodopa and naltrexone had two computational effects—an increased learning rate for avoided pain, and an increased degree of choice stochasticity—whose combination yielded no significant effects on basic, model-independent, performance measures.

### fMRI analyses

Next, we will report two sets of fMRI analyses. First, we performed an axiomatic analysis to identify brain activation encoding *general* aversive prediction errors (i.e., activation encoding the degree to which both pain and no-pain outcomes are relatively worse, or less good, than expected). In our previous study, this analysis revealed a general aversive prediction error signal in the PAG (Roy et al. 2014). Here, we examined whether we could replicate this finding, and whether our pharmacological manipulations affected the brain activation associated with the prediction-error axioms. Second, to address the question whether learning from received and avoided pain is supported by two separate brain systems, we sought to identify brain activation encoding *outcome-specific* prediction error signals. In both analyses, we focused on the first second of the outcome period as this is when prediction errors are triggered.

We modeled drug effects using two second-level regressors (levodopa vs. placebo and naltrexone vs. placebo) in both analyses. All fMRI results are thresholded at *q <* 0.05, false discovery rate (FDR)-corrected for multiple comparisons across the whole brain (gray matter masked), unless otherwise stated (e.g., for visualization purposes). Unthresholded *t* maps can be found on https://neurovault.org/collections/RIVRRMAK/.

#### General (outcome-nonspecific) aversive and appetitive prediction error signals

Prediction-error related activation is often examined by regressing fMRI activity at outcome onset on model-derived prediction errors (see Figure 3—figure supplement 1 for the corresponding activation in our study). However, as prediction errors are defined as the outcome minus the expected outcome, a problem with this approach is that the resulting brain activity may predominantly track the outcome (in our task: pain vs. no pain) or the expected outcome (in our task: the expected pain probability), which are intrinsically correlated with the prediction error.

**Figure 3.**
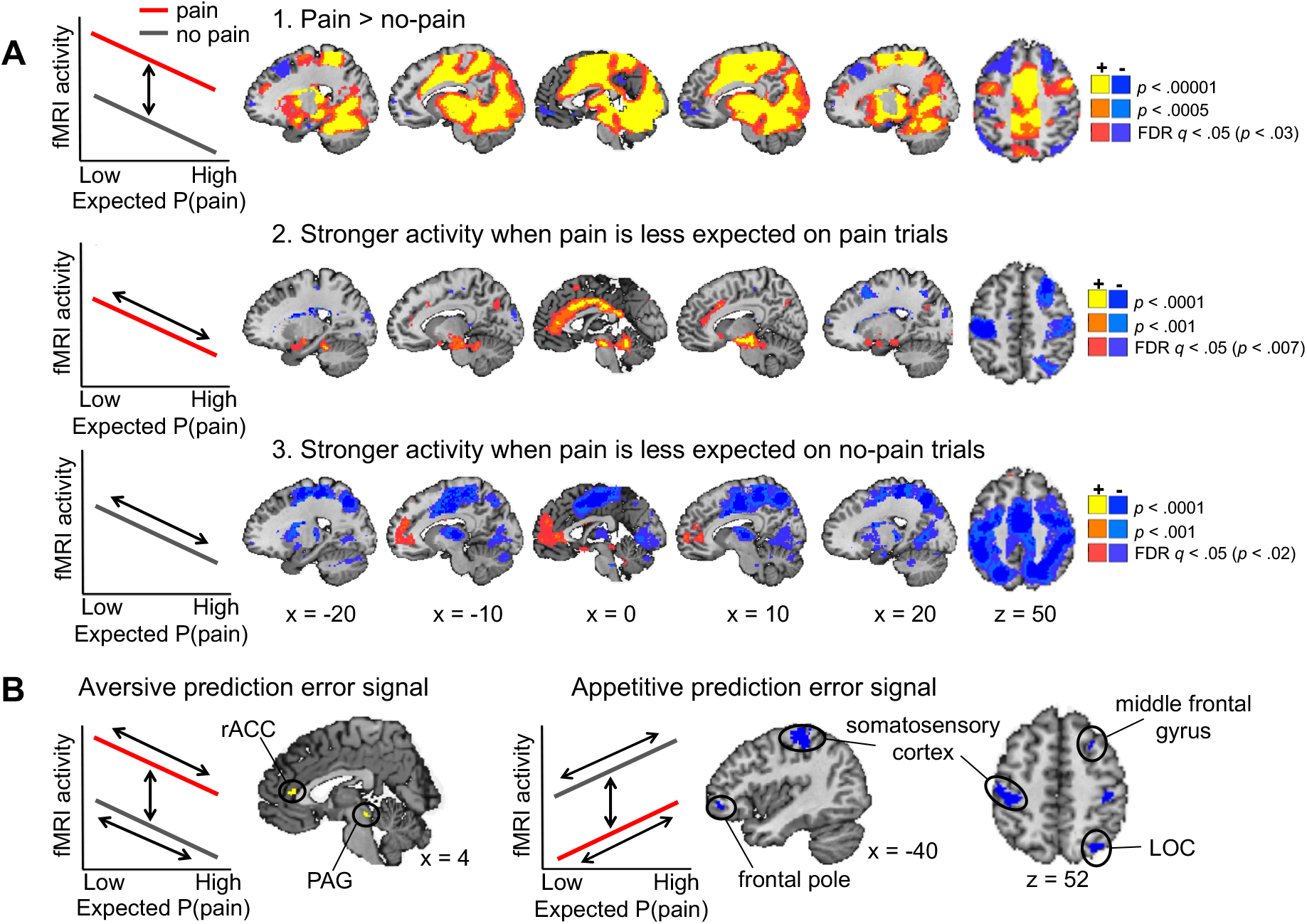
Axiomatic tests of brain activation encoding general aversive and appetitive prediction errors (N = 74). **A**. Activation associated with the three axioms for aversive prediction errors in our task. Yellow regions showed the effects illustrated in the left panels, and blue regions showed the reverse effects (i.e., the axioms for appetitive prediction errors). Expected P(pain) is expected pain probability. All maps were thresholded at *q <* 0.05, FDR-corrected for multiple comparisons across the whole brain, with higher voxel thresholds superimposed for display. **B**. Conjunction results. Regions activated for each of the above three contrasts, all thresholded at *q <* 0.05 FDR-corrected. Yellow and blue regions showed positive and negative responses for each contrast, respectively, thus encoded general aversive and appetitive prediction errors.

To address this issue, and identify brain activity that truly integrates actual and expected outcomes into a prediction error signal, a set of conditions has recently been specified (Rutledge et al. 2010, Roy et al. 2014). These conditions, or axioms, for general aversive prediction error signals in our task are: (i) activation at outcome onset should be higher for received than avoided pain, unless pain is fully expected; (ii) when pain is received, activation should be higher when pain was less expected (i.e., negative correlation with expected pain probability); and (iii) when pain is avoided, activation should also be higher when pain was less expected, that is, when avoidance was more expected (Figure 3A, left panels). To identify regions encoding a general aversive prediction-error signal, we tested for activation that fulfilled each of these three axioms, using a whole-brain conjunction analysis. In addition, to search for regions encoding the opposite (i.e., appetitive-like) prediction error signal, we also tested for activation that fulfilled each of the *reverse* axioms. Note that we did not detect activation encoding appetitive-like prediction errors in our previous study (Roy et al. 2014).

A fourth axiom, that applies to both aversive and appetitive prediction errors, is that activation for received and avoided pain should be equivalent if the outcome is fully predicted (i.e., when the prediction error is zero). As outcomes could never be fully predicted in our task, we could not test this axiom.

##### Axiom 1

A large part of the brain fulfilled the first axiom for aversive prediction errors (stronger response to received than avoided pain), including typical pain-processing regions such as the dorsal anterior cingulate cortex (ACC), (pre)motor cortex, insula and thalamus, as well as occipital (visual) cortex (Figure 3A, upper panel). We found the opposite effect (stronger response to avoided than received pain) in regions of the ventromedial prefrontal cortex (vmPFC), dorsolateral prefrontal cortex (dlPFC), somatosensory cortex, posterior ACC, and later occipital cortex (LOC).

##### Axiom 2

A test of the second axiom for aversive prediction errors (stronger responses to more unexpected pain) revealed several activation clusters, including regions in the vmPFC, dorsal ACC, insula, amygdala, and a midbrain area covering part of the PAG (Figure 3A, middle panel). In addition, several other regions, including the right dlPFC and bilateral somatosensory cortex, showed the opposite effect (stronger responses to more expected pain).

##### Axiom 3

The third axiom for aversive prediction errors (stronger responses to more expected pain avoidance) was fulfilled by a few regions in the vmPFC and rostral ACC (rACC), as well as part of the PAG. We also found the opposite effect (stronger responses to more unexpected pain avoidance) in several regions, including the dorsal ACC, sensorimotor cortex, thalamus, putamen and insula.

##### Conjunction

A conjunction analysis of the three contrasts reported above (Figure 3B) revealed two brain regions that satisfied all three axioms for aversive prediction errors: A midbrain region including part of the PAG (16 voxels) and an area in the rostral ACC (24 voxels). Importantly, we also identified activation that showed a *negative* effect for all three axioms—thus encoding appetitive-like prediction errors—in bilateral somatosensory cortex (433 and 65 voxels in the left and right hemisphere, respectively), left frontopolar cortex (47 voxels), right dlPFC (middle frontal gyrus; 27 voxels), and right LOC (253 voxels).

Together, these results replicate our previous finding that the PAG encodes general aversive prediction errors (Roy et al. 2014), and suggest a role for the rostral ACC in encoding aversive prediction errors as well. Furthermore, the identification of an additional neural circuit encoding appetitive-like prediction errors provides an important extension to our previous results, possibly owing to the larger number of participants and hence higher power in the present study.

#### No drug effects on prediction-error related brain activation

We found no differences between the levodopa and placebo group, nor between the naltrexone and placebo group, for any of the three prediction-error contrasts reported above (whole-brain FDR-corrected). Drug effects were virtually absent at lower, uncorrected, thresholds as well (see unthresholded *t* maps on https://neurovault.org/collections/RIVRRMAK/). More specific tests for drug effects in the clusters identified by our conjunction analysis provided no evidence for effects of our pharmacological manipulations on prediction-error related brain activation either (Figure 3— figure supplement 2).

#### Outcome-specific prediction error signals

The previous analysis identified regions that encoded the degree to which outcomes were relatively worse- or better-than-expected in the *same* direction when pain was received and avoided, but did not *dissociate* learning from received and avoided pain. To address this issue, we next examined whether different brain regions encode the unexpectedness, or surprise (which drives learning in reinforcement-learning models), evoked by received and avoided pain.

Regions encoding the surprise evoked by received pain should respond stronger to pain outcomes when pain was less expected (i.e., a negative correlation with expected pain probability on pain trials). In contrast, regions encoding the surprise evoked by avoided pain should respond stronger to no-pain outcomes when pain was more expected (i.e., a positive correlation with expected pain probability on no-pain trials; Figure 4A, left plots). To identify regions that show a stronger surprise response for received than avoided pain, we thus specified the following contrast: negative correlation with expected pain probability on pain trials > positive correlation with expected pain probability on no-pain trials. This contrast revealed activation in the vmPFC and rACC, posterior cingulate cortex, insula extending into the left parahippocampal gyrus, cerebellum and a brainstem region encompassing part of the PAG (Figure 4A, yellow regions). We also found extensive clusters that showed the opposite effect—i.e. positive correlation with expected pain probability on no-pain trials > negative correlation with expected pain probability on pain trials—in sensorimotor cortex, parietal and occipital cortex, and the bilateral frontal poles (Figure 4A, blue regions), suggesting that these regions show a stronger surprise response for avoided than received pain.

**Figure 4.**
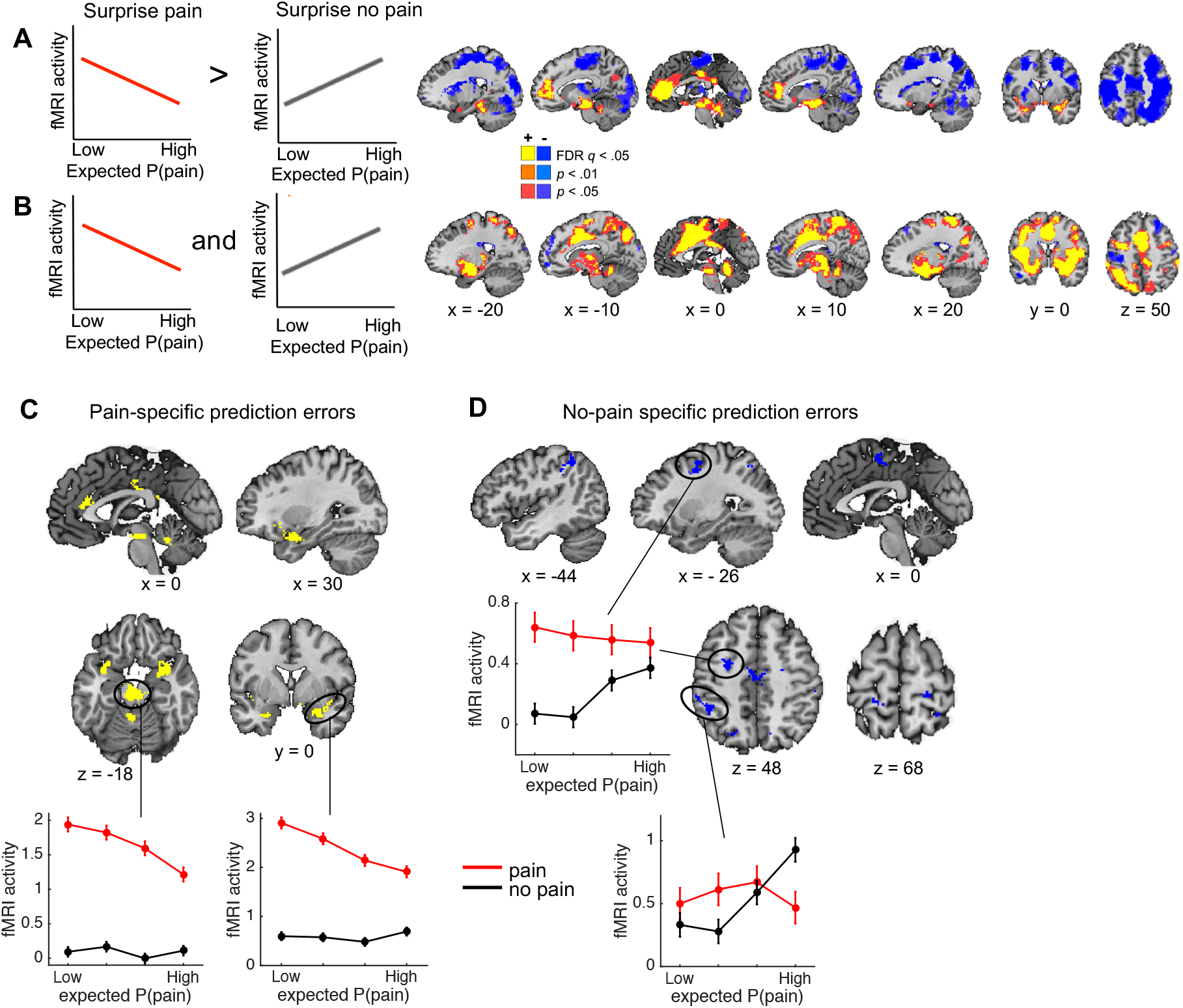
Outcome-specific prediction-error signals (N = 74). **A**. Activation tracking surprise more for received than avoided pain (yellow) and vice versa (blue). Note that this includes activation that tracks expected pain probability across both outcomes. **B**. Activation tracking surprise for both received and avoided pain (i.e., absolute prediction error). Activation maps in A and B are thresholded at *q <* 0.05, FDR-corrected for multiple comparisons across the whole brain, with adjacent areas thresholded at p < 0.01 and p < 0.05 (uncorrected) for display. **C**. Regions encoding surprise more for received than avoided pain, which cannot be explained by a general sensitivity to expected pain probability. These regions showed positive activation for both the first (A) and second (B) contrast, each thresholded at *q <* 0.05, FDR-corrected. **D**. Regions encoding surprise more for avoided than received pain, which cannot be explained by a general sensitivity to expected pain probability. These regions showed negative activation for the first, and positive activation for the second contrast, each thresholded at *q <* 0.05, FDR-corrected. The line plots show the mean activity extracted from the brainstem and right amygdala (C) and left dlPFC and parietal (D) clusters per quartile of expected pain probability, illustrating the encoding of outcome-specific prediction errors in these regions.

We also sought to identify activation encoding the surprise elicited by both received and avoided pain, that is, activation encoding absolute prediction errors. To this end, we specified a second contrast that tested for a negative correlation with expected pain probability on pain trials *and* a positive correlation with expected pain probability on no-pain trials (Figure 4B). This contrast revealed extensive activation clusters in the dorsal ACC extending into the supplementary motor cortex, insula, sensorimotor cortex, thalamus, part of the brainstem and cerebellum, suggesting that these regions encoded absolute prediction error (Figure 4B, yellow regions). In addition, a few smaller clusters in left sensorimotor cortex, right dlPFC, and left frontopolar cortex showed the opposite effect, suggesting that these regions encoded the overall *expectedness* of outcomes (Figure 4B, blue regions).

Finally, note that a caveat of the first contrast reported above (Figure 4A) is that it also identifies activation that is stronger when the expected pain probability is lower (i.e., activation encoding expected safety), regardless of the outcome. This likely explains the vmPFC activation, which has been found to represent positive affective value in many domains (Roy et al. 2012, Bartra et al. 2013, Rich and Wallis 2016). The second contrast (for absolute prediction errors; Figure 4B), on the other hand, will *not* detect activation encoding expected safety regardless of the outcome, as the correlation with expected pain probability is specified in opposite directions for pain and no-pain outcomes. Thus, we reasoned that regions identified by both of the contrasts reported in Figure 4A and 4B encode outcome-specific prediction errors (Figure 4A) unconfounded by outcome-nonspecific expected pain probability (Figure 4B). Therefore, we next examined the conjunction of these two contrasts. Specifically, we masked the activation identified by the first contrast (separately for the positive and negative activation) by the positive activation identified by the second contrast. Positive activation for both contrasts was found in a set of mostly subcortical and limbic regions, including a large cluster in the brainstem (not covering the PAG), bilateral insula extending into the amygdala, rostral ACC, posterior cingulate cortex, bilateral supramarginal gyrus, and cerebellum (Figure 4C). These regions thus encoded surprise more for received than avoided pain (pain-specific prediction errors), which could not be explained by a general sensitivity to expected pain probability. Negative activation for the first contrast and positive activation for the second contrast, on the other hand, was found in several cortical regions, including the supplementary motor cortex, left parietal cortex (supramarginal gyrus), bilateral somatosensory cortex (postcentral gyrus), and left dlPFC (middle and superior frontal gyrus) (Figure 4D). These regions thus encoded surprise more for avoided than received pain (no-pain specific prediction errors), which could not be explained by a general sensitivity to expected pain probability. Together, these results provide evidence that prediction errors evoked by pain and no-pain outcomes are encoded in largely distinct brain regions.

#### No drug effects on surprise-related brain activation

Because levodopa and naltrexone specifically increased learning rates for no-pain outcomes, we expected these drugs to increase prediction-error related brain activation for no-pain outcomes as well. However, we found no differences between the levodopa and placebo group, or between the naltrexone and placebo group, for any of the contrasts reported above (whole-brain FDR-corrected). Drug effects were virtually absent at lower, uncorrected, thresholds as well (see unthresholded *t* maps on https://neurovault.org/collections/RIVRRMAK/).

## Discussion

Our results provide novel evidence that unexpectedly received and avoided pain— signaling threat and safety, respectively—drive human pain-avoidance learning via different learning systems. First, computational modeling suggested that participants’ choices were best explained by a model with separate learning rates for received and avoided pain, and that untreated participants learned more from received than avoided pain. Second, levodopa and naltrexone selectively increased learning rates for avoided pain, suggesting a role for the dopamine and endogenous-opioid systems in safety, but not threat, learning. Third, our fMRI analyses revealed that different brain circuits encode prediction errors elicited by received and avoided pain, providing evidence for two dissociable learning systems at the neural level as well. Somewhat surprisingly, however, we found no drug effects on fMRI activity for any of the contrasts we examined. We discuss each of these findings below.

### Learning rates for received vs. successfully avoided pain

The higher learning rate for pain than no-pain outcomes in the placebo group suggests that people normally (in the absence of a pharmacological manipulation) update their expectations more following received than avoided pain. A similar learning-rate asymmetry was found in a recent aversive reversal-learning study (Wise et al. 2019). Interestingly, however, reward-learning studies using secondary outcomes (e.g., monetary gains and losses) have provided evidence for the *opposite* asymmetry: higher learning rates for favorable than unfavorable outcomes. This has been attributed to an optimistic learning bias (Sharot and Garrett 2016, Lefebvre et al. 2017) and a tendency to learn preferentially from information that confirms one’s current action (Palminteri et al. 2017). The opposite learning bias in pain-avoidance learning tasks may be due to the intrinsically aversive nature of pain, arguably rendering unexpected pain a more salient teaching signal than unexpected pain absence. Relatedly, the experience of pain may trigger a reflexive tendency to change one’s course of action (Huys et al. 2012), expressed in elevated learning rates for pain outcomes. Thus, higher learning rates for received than avoided pain may reflect a Pavlovian influence on choice which operates in parallel to the instrumental learning system. Alternatively, the seemingly opposite learning asymmetries in reward-learning and pain-avoidance learning tasks may also reflect a cognitive process related to the framing of the task. That is, participants who are instructed to maximize reward vs. minimize pain may pay most attention to—and hence learn most from—reward vs. pain outcomes, respectively.

The presence and direction of learning asymmetries may also depend on the specific task demands. For example, a previous fMRI study that used a more complex pain-avoidance learning task (pain probabilities of three different options were learned in parallel, in an indirect manner), and included a risk-taking component, found no systematic difference in learning rates for received and avoided pain (Eldar et al. 2016). Interestingly, in contrast to our present and previous (Roy et al. 2014) fMRI findings, the PAG in that study *positively* encoded expected pain probability on no-pain trials (one of the axioms for appetitive prediction errors), and did not encode expected pain probability on pain trials. These findings suggest that the task used in that previous study (Eldar et al. 2016) evoked different learning processes than our simpler task. Indeed, learning rates in that previous study were much lower than those found in our task as well. Further examination of the degree to which avoidance-learning processes and their neural implementation generalize across different learning tasks is an important objective for future work (Yarkoni 2020).

### Effects of levodopa and naltrexone on learning parameters

Levodopa and naltrexone selectively increased learning rates for successfully avoided pain, consistent with a role for dopamine and endogenous opioids in safety learning. Previous studies have associated levodopa-induced increases in phasic dopamine activity with enhanced learning from (secondary) rewards (Frank et al. 2004, Pessiglione et al. 2006). Combined with these previous findings, our levodopa results suggest that phasic dopamine activity may signal the degree to which outcomes are ‘better than expected’ across both reward and punishment domains. We are aware of one previous study that provided correlational evidence for a role of dopamine in human safety learning in a Pavlovian fear-conditioning task (Raczka et al. 2011). That study found that individual differences in fear-extinction learning rates were associated with genetic variation in the dopamine transporter gene, which presumably affects phasic striatal dopamine release. Our levodopa results are consistent with this result, and provide the first *causal* evidence for a role of dopamine in human safety learning in an instrumental-learning task.

It has been proposed that the dopamine system signals aversive prediction errors as well, via a subset of midbrain dopaminergic neurons that is responsive to aversive outcomes (Brooks and Berns 2013). Consistent with this hypothesis, a subset of putative dopamine neurons in non-human primates (Matsumoto and Hikosaka 2009) and rodents (Brischoux et al. 2009, Lammel et al. 2011) have been found to increase their activity in response to aversive stimuli. However, it is yet unclear whether these neurons signal aversiveness, salience, or relief from a threat period (Mirenowicz and Schultz 1996, Schultz 2019), and whether these neurons are really dopaminergic (Ungless et al. 2004). In our study, levodopa did not affect learning rates for received pain; hence our results do not support the idea that dopamine supports learning from aversive outcomes. According to another account—which is largely based on studies using secondary reinforcers in Parkinson’s disease patients— punishment learning (e.g., learning from monetary losses) is mediated by phasic *decreases* (dips) or pauses in dopamine activity, via an effect on D2 receptors (Frank 2005, Maia and Frank 2011). Although levodopa is thought to primarily increase phasic dopamine bursts (Floel et al. 2008, Black et al. 2015), it may increase tonic dopamine levels as well, which could in turn interfere with phasic dopamine dips (Breitenstein et al. 2006, Moustafa et al. 2013, Poletti and Bonuccelli 2013). Thus, if dopamine dips following aversive prediction errors support pain-avoidance learning, we would expect levodopa to *impair* learning from received pain. Our finding that levodopa did not affect learning rates for received pain does not support this hypothesis either. Instead, our results suggest a selective role for the human dopamine system in learning from successfully avoided pain.

Regarding the endogenous opioid system, we expected that if pain-avoidance learning relies on μ-opioid activity, naltrexone—which blocks this activity—would *suppress* learning from received and/or avoided pain. However, we found the opposite effect for avoided-pain outcomes: like levodopa, naltrexone increased learning rates for avoided pain. This finding counterintuitively suggests that μ-opioid activity normally suppresses learning from avoided pain and that naltrexone countered this effect, which seems to contradict findings that μ-opioid receptor antagonists impair fear-extinction learning in rats (McNally and Westbrook 2003, McNally et al. 2004, McNally et al. 2005, Kim and Richardson 2009, Parsons et al. 2010). Obviously, these animal studies differed from our study in several ways—such as the nature of the learning task (Pavlovian vs. instrumental), outcome measure (freezing behavior vs. learning-rate estimates), opioid-receptor antagonist (naloxone vs. naltrexone) and drug administration (injection into the PAG vs. oral administration)—each of which may explain the apparently contradictory results. Thus, our results show that untreated participants learn more rapidly from received than avoided pain (replicated in our previous study) and that both levodopa and naltrexone negate this learning asymmetry, but the neurobiological mechanisms underlying the naltrexone effect remain to be elucidated. One informative approach for future studies would be to directly compare effects of opioid-receptor agonists and antagonists on pain-avoidance learning parameters.

The levodopa and naltrexone groups also showed a higher degree of choice stochasticity than the placebo group, suggesting that participants in both drug groups were less prone to choose the option with the lowest expected pain probability. Importantly, our parameter-recovery analysis indicated that the drug effects on learning rate for avoided pain and choice stochasticity could be independently and correctly retrieved. The increased choice stochasticity in the levodopa group may reflect a positive association between dopamine activity and risk preference (Voon et al. 2006, Gallagher et al. 2007, St Onge and Floresco 2009, Chew et al. 2019) or exploration (Beeler et al. 2010, Kayser et al. 2015, Gershman and Tzovaras 2018). It is also broadly consistent with recent evidence that levodopa reduces the impact of valence on information-seeking (Vellani et al. 2020). We are not aware of previous studies that associated the endogenous opioid system with choice stochasticity, risk-taking or exploration. It is possible that the similar effects of levodopa and naltrexone on choice stochasticity were due to common general side effects of both drugs on participants’ attention or motivation, which disrupted the decision-making process. However, we believe this is unlikely because (i) the drugs did not affect subjective state (alertness, calmness or contentment) and (ii) general side effects cannot easily explain the specific increase in learning rates for avoided pain.

Interestingly, the drug effects on learning rate for avoided pain and choice stochasticity were not accompanied by drug effects on basic performance measures (number of received pain outcomes, pain-switch or avoid-switch behavior). In a similar vein, previous studies have reported effects of pharmacological manipulations and dopaminergic genotype on model parameters despite a lack of significant effects on behavioral measures (Raczka et al. 2011, Chakroun et al. 2020), illustrating the added value of computational models. Our simulation results suggested that the drug effects on learning rate and choice stochasticity in our study cancelled each other out. Specifically, symmetric learning rates for received and avoided pain (found in the drug groups) result in more accurate pain-probability estimates than asymmetric learning rates (found in the placebo group). This beneficial effect of a symmetric learning process in the drug groups was, however, counteracted by the detrimental effect of a more stochastic choice process, resulting in no net performance difference between the placebo and drug groups.

### Separate brain circuits support learning from received and avoided pain

Our fMRI results provided evidence for two dissociable learning systems at the brain level as well. Pain-specific prediction errors were predominantly encoded in subcortical (brainstem, cerebellum) and limbic (insula, amygdala, rostral ACC) regions that are typically associated with emotional and affective processes, including fear conditioning (Phillips and LeDoux 1992) and affective responses to errors (Bush et al. 2000). In contrast, no-pain specific prediction errors were encoded in frontal and parietal cortical areas typically associated with higher-order cognitive processing (Ptak et al. 2017).

Prediction errors for no-pain outcomes were not represented in the ventral striatum, which is typically found for reward prediction errors (O’Doherty et al. 2003, Rutledge et al. 2010). Instead, prediction errors for no-pain outcomes were associated with frontoparietal activity. This activity is unlikely to reflect a reward signal, but possibly reflected increased attention on trials in which pain was expected but not received. That is, the unexpected absence of pain may have prompted participants to carefully monitor the thermode’s temperature in order to verify whether pain was really avoided or still to come, as reflected in increased frontoparietal activity. The absence of a striatal ‘reward-like’ prediction-error response for no-pain outcomes suggests that, in terms of neural processing, avoiding pain is not comparable to gaining a reward. However, the lack of a detectable reward-like prediction-error response may also be related to our task design. Specifically, pain outcomes in our task involved a change in sensory input (a rise in temperature) whereas no-pain outcomes did not (maintenance of the baseline temperature), which may have caused a more prominent neural prediction-error response for the pain outcomes. One way to examine this issue would be to use a task in which choices result in either an increase or a decrease in painful stimulus intensity from a tonic pain level, such that aversive and appetitive-like outcomes are associated with similar changes in sensory input (Seymour et al. 2005). Such a task would examine pain-relief rather than pain-avoidance learning. The current task design, however, more closely resembles the situation of a patient recovering from injury or surgery, who is pain-free as long as he or she is resting but expects that physical activity may result in pain. It is an interesting speculation that, in such situations, the stronger subcortical and limbic (’emotional’) responses for pain than for no-pain prediction errors, as found in our study, may foster a behavioral state that favors inactivity and rest, which could promote tissue healing and recovery.

### No effects of levodopa and naltrexone on fMRI activation

Unexpectedly, we found no effects of levodopa or naltrexone on any of our fMRI prediction-error contrasts. This may indicate that the dopamine and opioid systems are not involved in pain-avoidance learning, although this is inconsistent with the drug effects on learning rates for avoided pain. It is also possible that our pharmacological manipulations *did* affect prediction-error related dopamine and/or opioid activity, but that we did not have enough power to detect these effects due to our moderate sample size and between-subject design (24-26 participants per treatment group). Relatedly, drug effects on prediction-error related dopamine and/or opioid activity may not always produce corresponding changes in the BOLD signal (Knutson and Gibbs 2007, Brocka et al. 2018). This last possibility is clearly discouraging for pharmacological fMRI studies, but is conceivable given the absence of consistent levodopa effects on reward prediction-error signals in previous fMRI studies: One study reported an increased reward prediction-error signal in an levodopa group compared to a haloperidol group (although neither of these groups differed from a placebo group) (Pessiglione et al. 2006), but two recent studies using within-subject designs found no levodopa effects on neural reward prediction-error signals (Kroemer et al. 2019, Chakroun et al. 2020). To better understand the degree to which drug-induced changes in human neuromodulatory (e.g., dopamine) activity are detectable using fMRI, future research could directly compare drug effects on local changes in neuromodulator activity—e.g., using molecular imaging procedures such as PET—with effects of these same drugs on the BOLD signal.

### Limitations and directions for future research

As mentioned above, a lack of statistical power due to our moderate sample size and between-subject design may have prevented the detection of drug effects in our fMRI analyses. In addition, our fMRI data suffered from signal dropout in inferior parts of the prefrontal cortex (including the orbitofrontal cortex), which is a common problem in fMRI studies using echo-planar imaging (Ojemann et al. 1997, Deichmann et al. 2003). Therefore, our results are agnostic with respect to the contribution of ventral prefrontal areas to pain-avoidance learning, and their potential modulation by levodopa or naltrexone. Finally, regarding the role of neuromodulators, we focused on dopamine and endogenous opioids, but other neuromodulators are almost certainly involved in pain-avoidance learning as well. In particular, future work may focus on the serotonergic system, which has traditionally been associated with aversive processing, behavioral inhibition, and “fight or flight” responses in rodents (Deakin 1983, Soubrie 1986, Deakin and Graeff 1991). More recent pharmacological and genetic studies, which mostly used tasks with secondary reinforcers, have provided evidence for a role of the human serotonergic system in various aspects of aversive learning (Chamberlain et al. 2006, Cools et al. 2008, Crockett et al. 2012, Hindi Attar et al. 2012, Robinson et al. 2012, den Ouden et al. 2013). Based on these findings, it has been proposed that serotonin acts as an opponent to dopamine by mediating behavioral inhibition in response to punishment (Daw et al. 2002, Dayan and Huys 2009, Boureau and Dayan 2011, Cools et al. 2011). When generalizing these findings to the domain of pain-avoidance learning, we may expect a specific role for the serotonergic system learning from received pain (threat learning). However, other studies have found effects of serotonin manipulations on both reward and punishment learning (Palminteri et al. 2012, Guitart-Masip et al. 2014), and a study in which participants simultaneously learned probabilities of monetary rewards and painful shocks suggested that serotonin selectively modulates reward processing (Seymour et al. 2012). Taken together, previous work suggests that the serotonergic system has highly intricate and multifaceted roles in affective learning, likely due to its large number of receptor types, widespread projections, and interactions with other neuromodulators (Dayan and Huys 2009). Future studies are required to further clarify the functional role(s) of serotonin in pain-avoidance learning.

## Conclusion

In sum, our results suggest that received and avoided pain drive human pain-avoidance learning via two different learning systems, in terms of both learning rates and the neural encoding of prediction errors. In addition, our computational-modeling results provide evidence for a causal role of the dopamine and endogenous opioid systems in learning from avoided, but not received, pain. Future studies are needed to elucidate the neural mechanisms via which our dopamine and opioid manipulations affected learning rates for avoided pain, and to reveal the potential role of other neuromodulators in pain-avoidance learning.

## Methods

### Participants

Ninety-one healthy students (18-26 years old; 71% female; all right-handed) took part in the study. Participants reported no history of psychiatric, neurological, or pain disorders, and no current pain. Participants were instructed to abstain from using alcohol or recreational drugs 24 hours prior to testing, and to not eat or drink (except for water) 2 hours prior to testing. The study was approved by the medical ethics committee of the Leiden University Medical Center, and all participants provided written informed consent. Participants received a fixed amount of €60 plus a variable bonus of maximally €5 related to their performance on an additional task.

Six participants were excluded from all analyses because of thermode failure, and two additional participants because of poor task performance (see ‘Pain-avoidance learning task’ section below). In addition, nine participants were excluded from the fMRI, but not the behavioral, analyses because of excessive head movement (> 3 mm in any direction). Thus, the final behavioral analyses included 83 participants (placebo group: N = 28, mean age = 21.2, 74% female; levodopa group: N = 26, mean age = 20.8, 73% female; naltrexone group: N = 29, mean age = 20.8, 68% female). The final fMRI analyses included 74 participants (placebo group: N = 24, mean age = 21.3, 71% female; levodopa group: N = 24, mean age = 20.9, 71% female; naltrexone group: N = 26, mean age = 20.8, 69% female). Sample size was based on previous studies that detected effects of dopamine (levodopa) or opioid (naloxone) manipulations on behavioral and/or fMRI measures using between-subject designs (Pessiglione et al. 2006, Eippert et al. 2008, Guitart-Masip et al. 2012, Oei et al. 2012, Beierholm et al. 2013, Bunzeck et al. 2014, Guitart-Masip et al. 2014, Wittmann and D’Esposito 2015). These previous studies used sample sizes ranging from 13 to 30 participants per treatment group. We aimed at a sample size at the higher end of that range (25-30 participants per treatment group).

### General procedure

Two to fourteen days prior to the fMRI session, we assessed participants’ eligibility using a general health questionnaire and an fMRI safety screening form. During this screening session, participants also practiced the pain-avoidance learning task.

Each eligible participant took part in one fMRI session. On the day of the fMRI session, participants received a single oral dose of either 100 mg levodopa, 50 mg naltrexone, or placebo, according to a double-blind, randomized, between-subject design. Levodopa was combined with 25 mg carbidopa—a decarboxylase inhibitor that does not cross the blood brain barrier—to inhibit the conversion of levodopa to dopamine in the periphery. Approximately 30 minutes after drug administration, participants were positioned in the MRI scanner, after which we acquired a high-resolution structural scan. Approximately 53 minutes after drug administration, participants completed a 5-minute pain-rating task during which they received a series of (unavoidable) heat stimuli of varying temperatures and rated their experienced pain following each stimulus (Figure 1—figure supplement 2). This task was included to test for drug effects on subjective pain responses, and to select a painful yet tolerable temperature for each participant in the pain-avoidance learning task. Sixty minutes after drug administration, roughly corresponding with peak plasma concentrations of levodopa and naltrexone, participants started the pain-avoidance learning task (described below), which lasted approximately 45 minutes. Following the pain-avoidance learning task, participants performed an 8-minute probabilistic reward-learning task (not reported here).

We measured participants’ subjective state at the beginning (before drug administration) and end (two hours after drug administration) of the test session, by means of visual analogue scales measuring alertness, calmness and contentment (Bond, 1974) (Figure 1—figure supplement 1). Both subjective state measures were collected outside the scanner.

### Pain-avoidance learning task

This instrumental pain-avoidance learning task contained 144 trials, divided in 4 runs of 36 trials. On each trial, participants made a choice between two options (a diamond and a circle). Choosing each option was associated with a specific probability of receiving a painful heat stimulation. The probabilities of receiving pain when choosing each option drifted across trials according to two independent random walks (Figure 1A). We used three different pairs of random walks (each pair crossed at least once); each pair was administered to approximately one third of the participants in each treatment group.

Each trial started with the presentation of the two choice options, randomly displayed at the left and right side of the screen for 1800 ms (Figure 1B). During this period, participants had to select one option by pressing a left or right button of the response unit, using their right index or middle finger, respectively. If participants did not respond in time (1.2 % of trials), the computer randomly selected an option for them. The chosen option was highlighted for 200 ms, followed by an anticipation period of 3, 5, or 7 seconds during which a white asterisk (*) was presented in the center of the screen. Then the outcome—a painful heat stimulus applied to participant’s leg for 1.9 s (see Thermal stimulation section below) or no stimulus—was presented. Outcome onset was accompanied by a change of the central asterisk to a colored plus sign (+). The plus sign was red or green during the first 200 ms of each pain and no-pain outcome, respectively, after which it turned white for the remainder of the outcome period. The color change was meant to prevent outcome uncertainty during the initial phase of the outcome period. Each trial ended with an inter-trial interval of 6, 8, or 10 seconds during which an asterisk was presented. Except for the outcome probabilities, participants were fully informed about the task structure and procedure.

During the fMRI session, one participant switched choices more frequently following the absence of pain than following pain, and one other participant did deliberately not make a choice on 17% of the trials, due to the use of irrelevant strategies. We excluded these participants from further analysis.

#### Thermal stimulation

Heat stimuli (ramp rate = 40°C/s; 1 second at target temperature) were applied to the inner side of participants’ left lower leg using a Contact Heat-Evoked Potential Stimulator (CHEPS; 27-mm diameter Peltier thermode; Medoc Ltd., Israel). After the initial pain-rating task, 16% of the participants (4 in the placebo, 4 in the levodopa, and 7 in the naltrexone group) indicated that they would not tolerate repeated stimulation at the highest temperature they had received so far (50°C). For those participants, we used a temperature of 49°C in the pain-avoidance learning task. For the remaining participants, we used a temperature of 50°C. Between stimulations the thermode maintained a baseline temperature of 32°C. The total duration of each stimulation was 1850 ms (425 ms ramp-up and ramp-down periods, 1 second at target temperature) for 49°C stimuli, and 1900 ms (450 ms ramp-up and ramp-down periods, 1 second at target temperature) for 50°C stimuli. After each scan run, we moved the thermode to a new site on the participant’s leg. To reduce the impact of potential site-specific habituation (Jepma et al. 2014), we administered one initial heat stimulus before starting the first trial on a new skin site.

### Behavioral analyses

We tested whether the total number of received pain stimuli, the proportion of pain trials followed by a switch to the other choice option, and the proportion of no-pain trials followed by a switch to the other choice option differed between the three treatment groups, using one-way ANOVAs. In addition, for each treatment group, we used logistic regression to analyze the probability of switching choices as a function of outcome (pain vs. no-pain) over the six previous trials.

### Computational modeling

We fitted two reinforcement learning (Q learning) models to participants’ choice data: one model with a single learning rate (Model 1), and one model with separate learning rates for pain and no-pain outcomes (Model 2). On each trial *t*, both models update the expected probability of pain for the selected stimulus *s*, *Q*_*s*_, in response to the experienced outcome *O* (pain or no pain), according to:

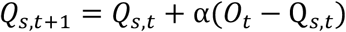

where (*O*_*t*_ − Q_*s,t*_) reflects the prediction error on each trial. We coded *O* as −1 and 0 for pain and no-pain outcomes, respectively, and initialized the *Q* values of both stimuli to −0.5. The value of the unchosen stimulus is not updated. Learning rate parameter α controls *how much* Q values are updated in response to each new outcome, such that higher values of α result in faster updating. The only difference between our two models is that Model 1 uses a single learning rate α for all outcomes, whereas Model 2 uses separate learning rates for pain and no-pain outcomes: *α*_*pain*_ and *α*_*no-pain*_, respectively.

Both models were combined with a softmax decision function, which computes the probability of choosing stimulus *s* on trial *t* (*P*_*s,t*_) as:

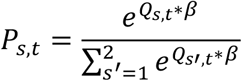

Inverse-temperature parameter *β* controls the sensitivity of choice probabilities to differences in Q values. If *β* is 0, both stimuli are equally likely to be chosen, irrespective of their expected pain probabilities. As *β* increases, the probability that the model chooses the stimulus with the lower expected pain probability increases. Thus, Model 1 has two free parameters (*α* and *β*) and Model 2 has three free parameters (*α*_*pain*_, *α*_*no-pain*_ and *β*).

#### Parameter estimation

We estimated the model parameters with a hierarchical Bayesian approach, using the hBayesDM package (Ahn et al. 2017). The hierarchical Bayesian approach assumes that every participant has a different set of model parameters, which are drawn from group-level prior distributions (Gelman 2014). The parameters governing the group-level prior distributions (hyperparameters) are also assigned prior distributions (hyperpriors). We estimated separate group-level parameters for the placebo, levodopa, and naltrexone groups. To test for treatment effects, we compared the posterior distributions of the group-level means (i.e., the hyperparameters governing the means of the group-level distributions) for each drug group vs. the placebo group.

##### Prior distributions

The group-level distributions for all individual parameters were assumed to be normal distributions. As hyperpriors for the mean and standard deviation (SD) hyperparameters of these group-level distributions we used, respectively, a normal distribution with mean 0 and SD 1 and a positive half-Cauchy distribution with location 0 and scale 5. We transformed the parameters’ unconstrained values to a [0,1] range using the inverse probit transformation (Ahn et al. 2017). In addition, we transformed *β* to a [0,20] range by multiplying its inverse-probit transformed values by 20.

##### MCMC sampling

The hBayesDM package performs hierarchical Bayesian parameter estimation with a Markov chain Monte Carlo (MCMC) sampling algorithm called Hamiltonian Monte Carlo (HMC) implemented in Stan and its R instantiation RStan. We ran 4 independent MCMC chains with different starting values, which each generated 5,000 posterior samples. We discarded the first 1,000 iterations of each chain as burn-in. In addition, we only used every 5th iteration to remove autocorrelation. Consequently, we obtained 3,200 representative samples (800 per chain) per parameter per model fit. All chains converged (Rhat values < 1.1).

#### Model comparison

We compared the fit of our two models using the Watanabe-Akaike information criterion (WAIC) (Watanabe 2010). The WAIC provides an estimate of a model’s out-of-sample predictive accuracy, adjusted for the number of free parameters, in a fully Bayesian way. Lower WAIC values indicate better out-of-sample predictive accuracy. WAIC is reported on the deviance scale (Gelman et al. 2014) hence a difference in WAIC value of 2 to 6 is considered positive evidence, a difference of 6 to 10 strong evidence, and a difference >10 very strong evidence for one model over another (Kass and Raftery 1995).

#### Computation of expected pain probability for use in fMRI analyses

We computed trial-specific expected pain probabilities—to be used as parametric modulator regressors in the fMRI analyses—by applying our winning model to each participant’s sequence of choices and outcomes. In line with previous studies (Pine et al. 2010, Voon et al. 2010, Seymour et al. 2012, Hauser et al. 2015, Kroemer et al. 2019), we used the exact same model to generate fMRI regressors for all participants by instantiating the model with the mean learning-rate parameters (the individual-level posterior medians, averaged across all participants).

### fMRI acquisition and analysis

#### Imaging acquisition

We acquired fMRI data on a 3T Philips Achieva MRI system (Best, The Netherlands) at the Leiden University Medical Center, using a standard whole-head coil. Stimulus presentation and data acquisition were controlled using E-Prime software (Psychology Software Tools). Visual stimuli were presented via a mirror attached to the head coil, and participants responded with their right hand via an MRI-compatible response unit. The pain-avoidance learning task was divided across four scan runs. Each run lasted 581 s (264 TRs), after discarding the first five TRs which served as dummy scans. Functional images were acquired with a T2*-weighted whole-brain echo-planar imaging sequence (TR = 2.2 s; TE = 30 ms, flip angle = 80°, 38 transverse slices oriented parallel to the anterior commissure-posterior commissure line, voxel size = 2.75 × 2.75 × 2.75 mm +10% interslice gap). In addition, we acquired a high-resolution T1-weighted scan (TR = 9.8 ms; TE = 4.6 ms, flip angle = 8°, 140 slices, 1.17 × 1.17 × 1.2 mm, FOV = 224×177×168), at the beginning of the scan session.

#### Preprocessing

Prior to preprocessing, global outlier time points (i.e. “spikes” in the BOLD signal) were identified by computing both the mean and the standard deviation (across voxels) of values for each image for all slices. Mahalanobis distances for the matrix of slice-wise mean and standard deviation values (concatenated) x functional volumes (time) were computed, and any values with a significant χ^2^ value (corrected for multiple comparisons based on the more stringent of either false discovery rate or Bonferroni methods) were considered outliers. On average 3.7% of images were outliers (SD = 1.9). The output of this procedure was later used as a covariate of noninterest in the first-level models.

Functional images were slice-acquisition-timing and motion corrected using SPM8 (Wellcome Trust Centre for Neuroimaging, London, UK). Structural T1-weighted images were coregistered to the first functional image for each subject using an iterative procedure of automated registration using mutual information coregistration in SPM8 and manual adjustment of the automated algorithm’s starting point until the automated procedure provided satisfactory alignment. Structural images were normalized to MNI space using SPM8, interpolated to 2×2×2 mm voxels, and smoothed using a 6mm full-width at half maximum Gaussian kernel.

#### Axiomatic analysis of general aversive and appetitive prediction error signals

For the first-level analysis, we create a general linear model (GLM) for each participant, concatenated over the four pain-avoidance learning blocks, in SPM8. We modeled periods of decision time (cue onset until response, mean response time = 758 ms), outcome anticipation (3-7 s), onsets of pain outcomes (1 s), and onsets of no-pain outcomes (1 s), using boxcar regressors convolved with the canonical hemodynamic response function. As in our previous study (Roy et al. 2014) we only modeled the first second of the outcome periods as this is when prediction errors are triggered. We added the model-derived expected pain probability as a parametric modulator on the outcome-anticipation and outcome-onset regressors. To control for potential effects of outcome-anticipation duration, we also included anticipation duration as a first parametric modulator on the outcome-onset regressors (using serial orthogonalization, such that any shared variance between expected pain probability and anticipation duration is assigned to the anticipation-duration effect). Other regressors of non-interest (nuisance variables) were i) “dummy” regressors coding for each run (intercept for each but the last run); ii) linear drift across time within each run; iii) the 6 estimated head movement parameters (x, y, z, roll, pitch, and yaw), their mean-zeroed squares, their derivatives, and squared derivatives for each run (total 24 columns per run); iv) indicator vectors for outlier time points identified based on their multivariate distance from the other images in the sample (see above); v) indicator vectors for the first two images in each run. Low-frequency noise was removed by employing a high-pass filter of 180 seconds.

For each participant, we created the following three contrast maps, corresponding to the axioms for general aversive prediction errors in our task: (i) pain onset > no-pain onset; (ii) negative correlation with expected pain probability at pain onset; and (iii) negative correlation with expected pain probability at no-pain onset. We performed a second-level (group) analysis on each of these contrasts using robust regression (Wager et al. 2005). We tested for drug effects by including two second-level regressors coding for levodopa vs. placebo (weights [-1 1 0] for the treatment groups [P L N]) and naltrexone vs. placebo (weights [-1 0 1] for the treatment groups [P L N]). All maps were thresholded at FDR *q* < 0.05 corrected for multiple comparisons across the whole brain (gray matter masked), with higher (more conservative) voxel thresholds superimposed for display in Figure 3A.

To identify brain regions that fulfilled all three axioms for general aversive prediction errors, we examined the conjunction between the three second-level contrast maps described above (pain > no pain, negative correlation with expected pain probability on pain trials, and negative correlation with expected pain probability on no-pain trials), each of them thresholded at FDR *q <* 0.05. To identify brain regions that fulfilled all three axioms for general appetitive prediction errors, we examined the conjunction between the three opposite contrast maps (no pain > pain, positive correlation with expected pain probability on pain trials, and positive correlation with expected pain probability on no-pain trials).

#### Analysis of outcome-specific prediction-error signals

To examine outcome-specific prediction-error signals, we used the same first-level GLM as in the previous analysis, but created different contrast maps. First, to identify activation that tracks surprise more for received than avoided pain (Figure 4A), we used the following contrast: ‘negative correlation with expected pain probability at pain onset’ > ‘positive correlation with expected pain probability at no-pain onset’. Second, to identify activation tracking absolute prediction error (Figure 4B), we used the following contrast: ‘negative correlation with expected pain probability at pain onset’ > ‘negative correlation with expected pain probability at no-pain onset’. Note that this contrast is identical to: ‘positive correlation with expected pain probability at no-pain onset’ > ‘positive correlation with expected pain probability at pain onset’. It is also identical to a contrast with weights [1 1] for the ‘negative correlation with expected pain probability at pain onset’ and positive correlation with expected pain probability at no-pain onset’ regressors.

We performed a second-level analysis on each of these two contrasts using robust regression, again including two second-level regressors coding for levodopa vs. placebo and naltrexone vs. placebo. Maps were again thresholded at FDR *q <* 0.05 corrected for multiple comparisons across the whole brain. Adjacent areas thresholded at p < 0.01 and p < 0.05 (uncorrected) were added for display in Figure 4A and 4B.

Finally, to identify regions encoding outcome-specific prediction errors which cannot be explained by a general sensitivity to expected pain probability (also see the Results section) we examined the conjunction between the two second-level contrast maps described above, each thresholded at FDR *q <* 0.05 (Figure 4C).

#### Conventional prediction error analysis

We also conducted a conventional (not axiomatic) analysis to identify aversive prediction-error related brain activation. In this analysis, we created a single regressor modeling all outcome onsets (pain and no pain) and added aversive prediction error (computed as the outcome [pain = 1, no pain = 0] minus the expected pain probability) as a parametric modulator on this outcome-onset regressor. The other regressors were identical to those in the previous analyses. We created first-level contrast images for the prediction error effect and conducted a second-level analysis on these contrast images, again thresholded at FDR *q <* 0.05 (Figure 3—figure supplement 1A). We also repeated this analysis while adding outcome (pain = 1, no pain = −1) as an additional parametric modulator on the outcome-onset regressor (without serial orthogonalization, such that only the unique variance explained by the prediction-error and outcome variables is assigned to their respective effects; Figure 3— figure supplement 1B).

## Acknowledgements

We thank Stephanie Bauduin, Laila Franke, Nikki Nibbering, Iliana Samara, and Iris Spruit for help with data collection.

## Competing interests

The authors declare that no competing interests exist.

## Data availability

Single-trial behavioral data and model code are available from the OSF database: https://osf.io/rqc6g/. Unthresholded *t* maps for all reported fMRI contrasts are available on Neurovault: https://neurovault.org/collections/RIVRRMAK/ (https://identifiers.org/neurovault.collection:6016).

---

**Figure 1 – Figure supplement 1.**
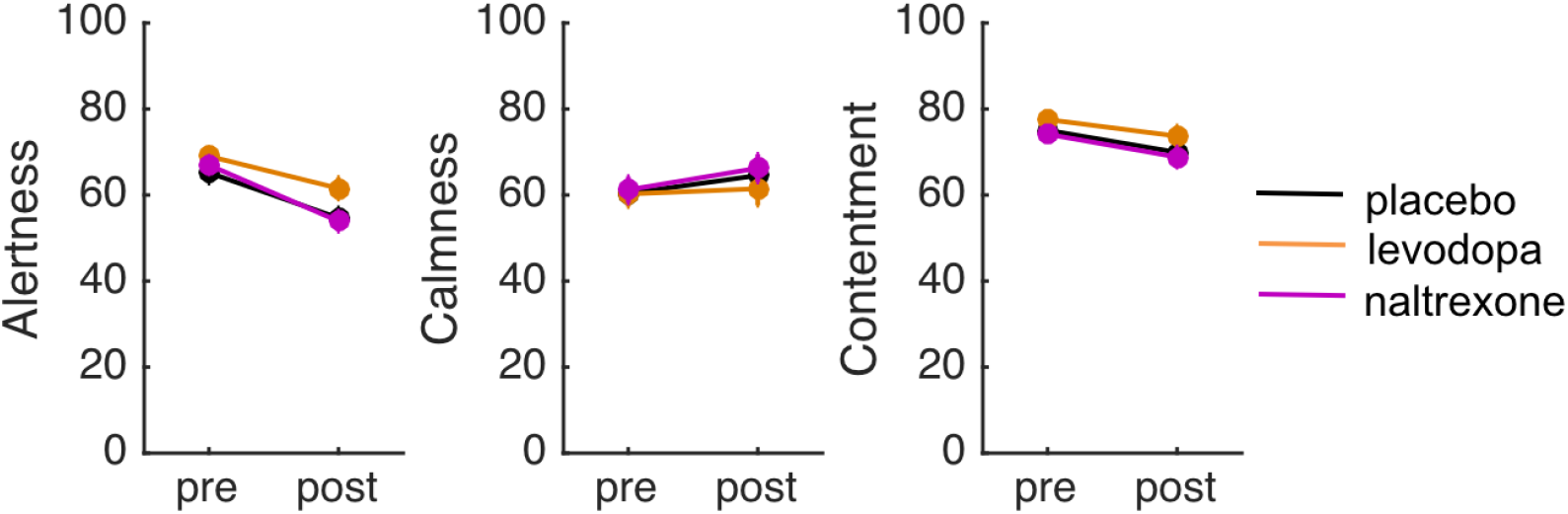
Pre- and post-treatment ratings of alertness, calmness, and contentment in each treatment group. Error bars indicate standard errors. The participants assigned to the three treatment groups did not differ in their pre-treatment ratings of alertness, calmness or contentment (all *p*’s>0.3). To assess potential treatment effects on subjective state we conducted analyses of covariance (ANCOVAs) on the post-treatment ratings of alertness, calmness and contentment (made two hours after drug intake), with treatment as a between-subject factor and the pre-treatment ratings as covariate. There was no effect of treatment on any of these ratings (all *p*’s>0.08), suggesting that the drugs did not affect subjective state.

**Figure 1 – Figure supplement 2.**
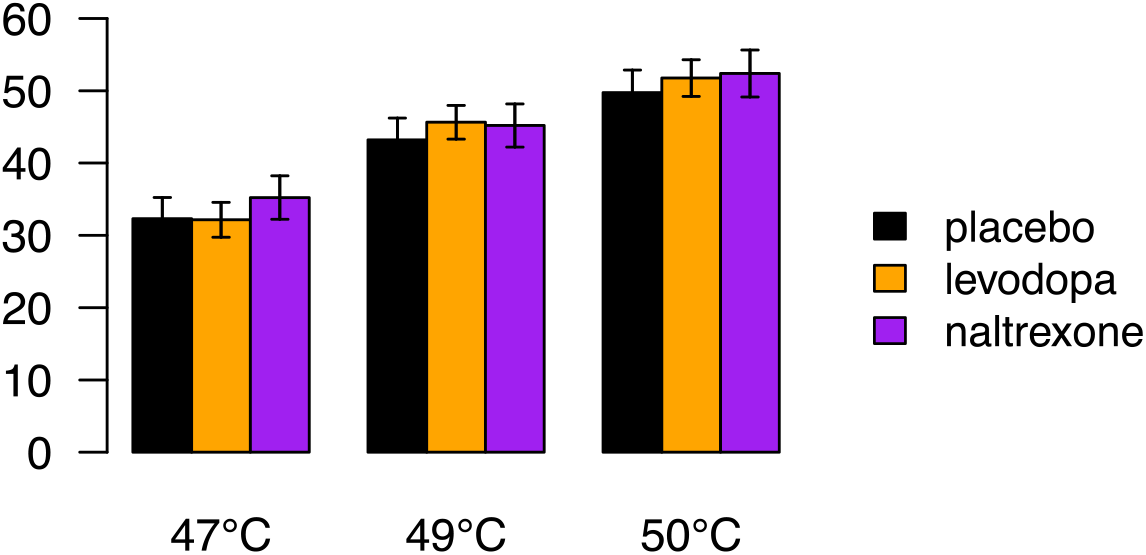
Pain ratings during a pain-rating task that preceded the pain-avoidance learning task, as a function of stimulus temperature and treatment group. Error bars indicate standard errors. Participants received five 47°C, five 49°C, and five 50°C heat stimuli, in random order, to their left lower leg (ramp rate = 40°C/second; 1 second at peak temperature, stimulus onset asynchrony = 17-25 seconds). Following each heat stimulus participants rated their experienced pain on a 100-unit visual analog scale with anchors of “no pain” and “worst-imaginable pain”, respectively. We conducted a mixed ANOVA on participants’ pain ratings with stimulus temperature (47°C, 49°C and 50°C coded as −1, 0 and 1, respectively) as within-subject factor and treatment as between-subject factor. Pain ratings increased as a function of stimulus temperature (*F*(1,84) = 424, *p <* 0.001). However, there was no main effect of treatment (*F*(2,84) = .21, *p =* .81) and no temperature x treatment interaction (*F*(2,84) = .76, *p =* .47). Thus, the drugs did not affect the subjective pain experience evoked by heat-pain stimuli. Note that four participants who were included in this analysis were excluded from the fMRI analyses of the pain-avoidance learning task, because of excessive head movement.

**Figure 2 – Figure supplement 1.**
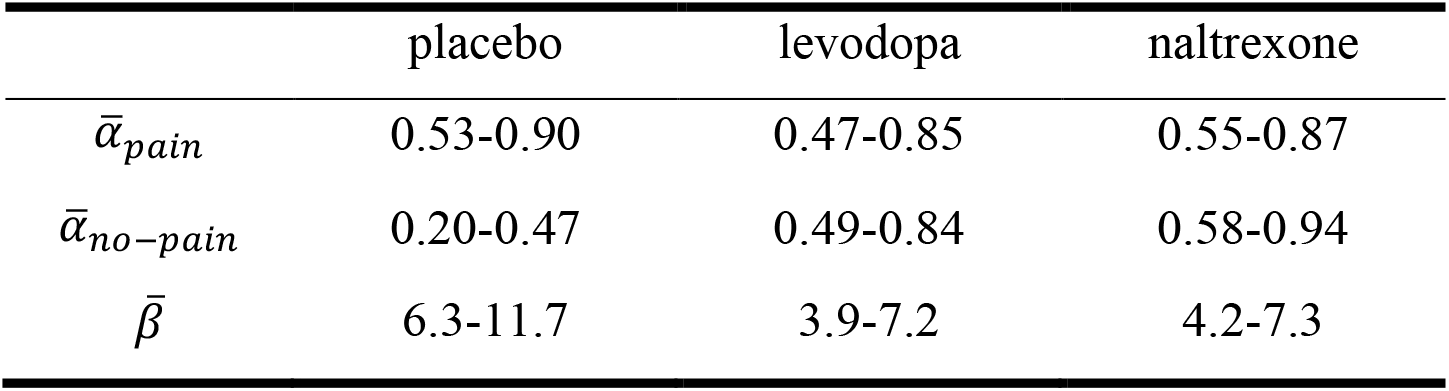
95% highest density intervals of the posterior distributions shown in Figure 2A.

**Figure 2 – Figure supplement 2.**
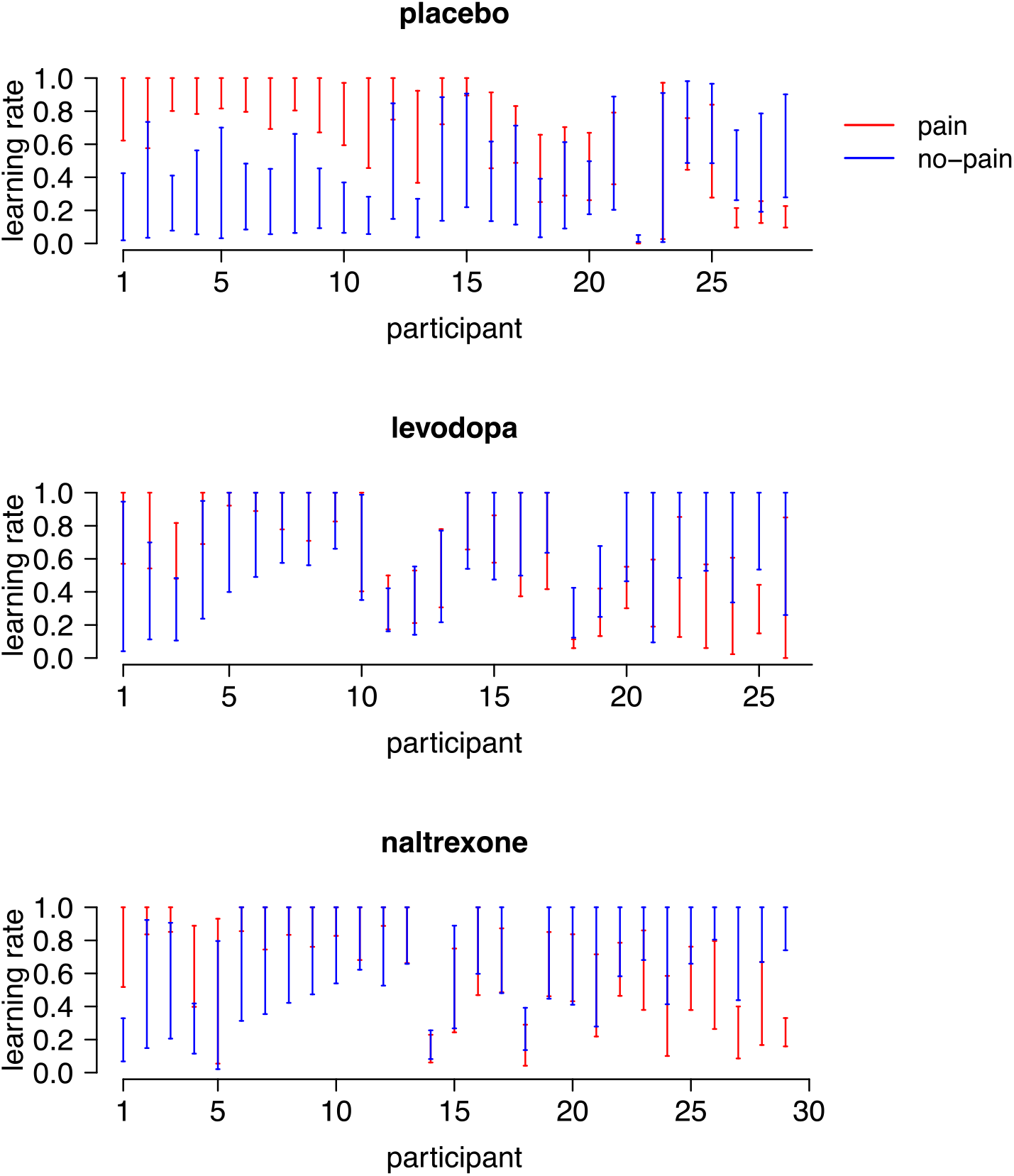
95% HDIs of the posterior distributions of each participant’s learning rate for pain (*α*_*pain*_) and no-pain (*α*_*no-pain*_) outcomes. Participants are sorted according to the difference between their two learning rates (*α*_*pain*_ − *α*_*no-pain*_). Note that *α*_*pain*_ and *α*_*no-pain*_ were positively correlated in the levodopa group (*r =* .43, *p =* .03), but were not correlated in the placebo (*r =* −.16, *p =* .4) and naltrexone (*r =* .31, *p =* .10) group.

**Figure 2 – Figure supplement 3.**
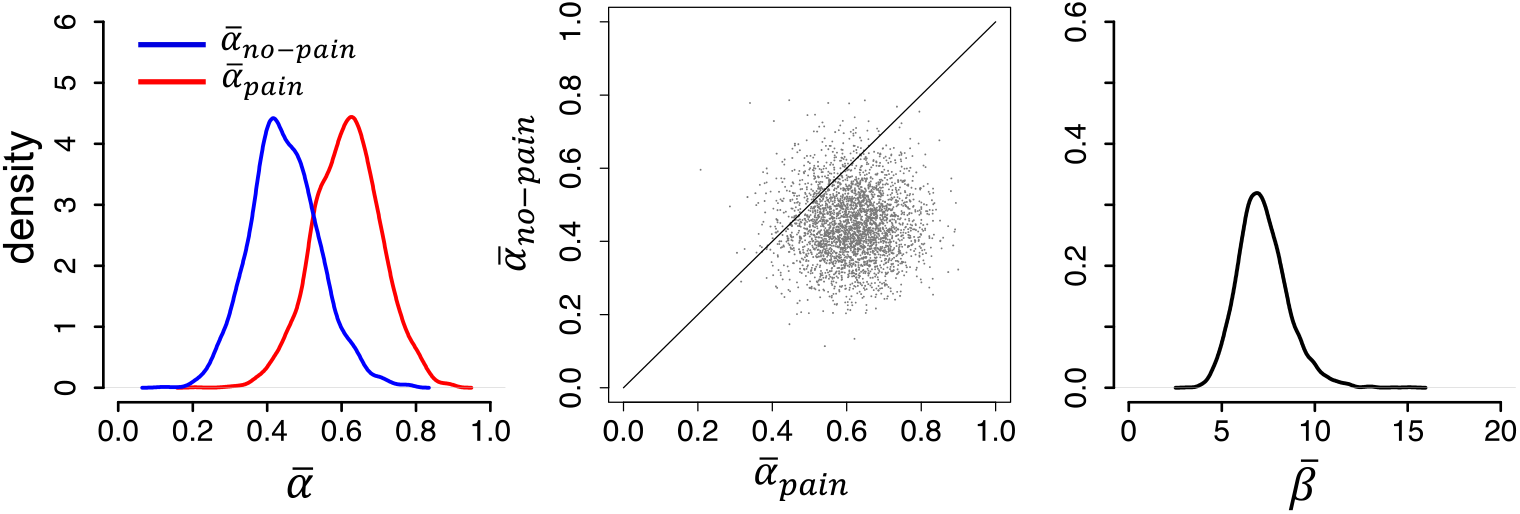
Modeling results from an independent sample of untreated participants from a previous study (N = 23), replicating the asymmetric learning rates (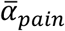 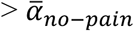) found in our placebo group.

**Figure 3 – Figure supplement 1.**
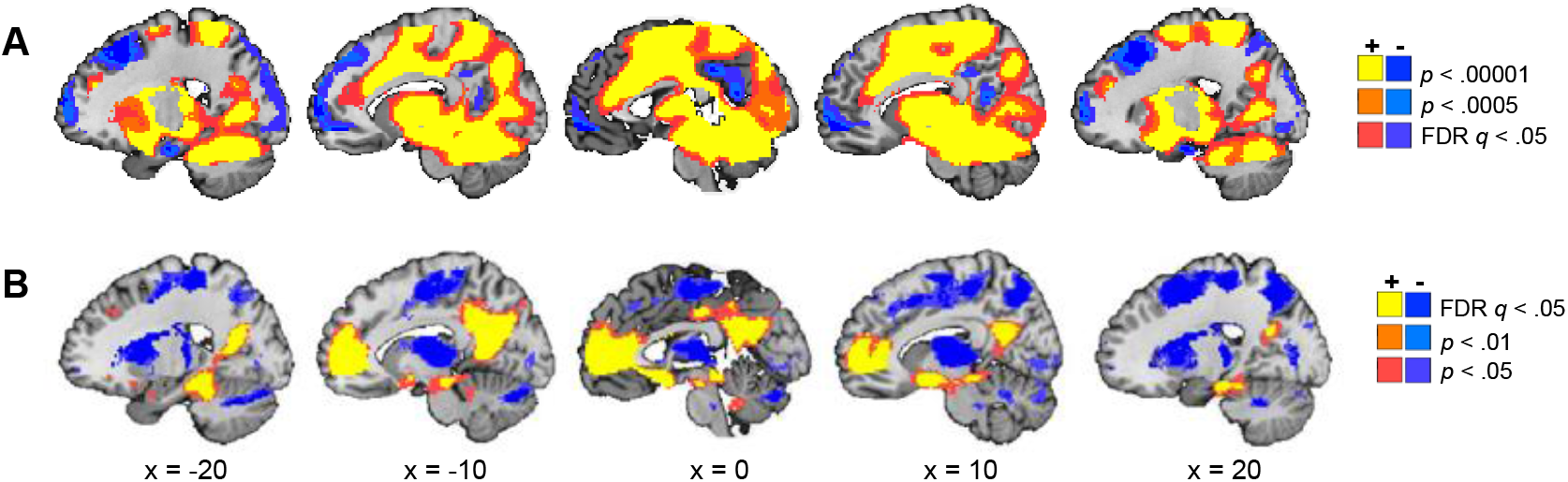
Results from a conventional prediction error analysis. **A**. Activation encoding general aversive prediction errors, defined as outcome (pain = 1, no pain = 0) minus model-derived expected pain probability. The trial-to-trial prediction error values were added as a parametric modulator on outcome onset. Note that the resulting activation resembles the activation found for the pain > no-pain contrast. **B**. Activation encoding general aversive prediction errors, controlling for outcome. Outcome (pain vs. no pain) was added as an additional parametric modulator on outcome onset. The resulting activation negatively correlates with expected pain probability (i.e., positively correlates with expected safety).

**Figure 3 – Figure supplement 2.**
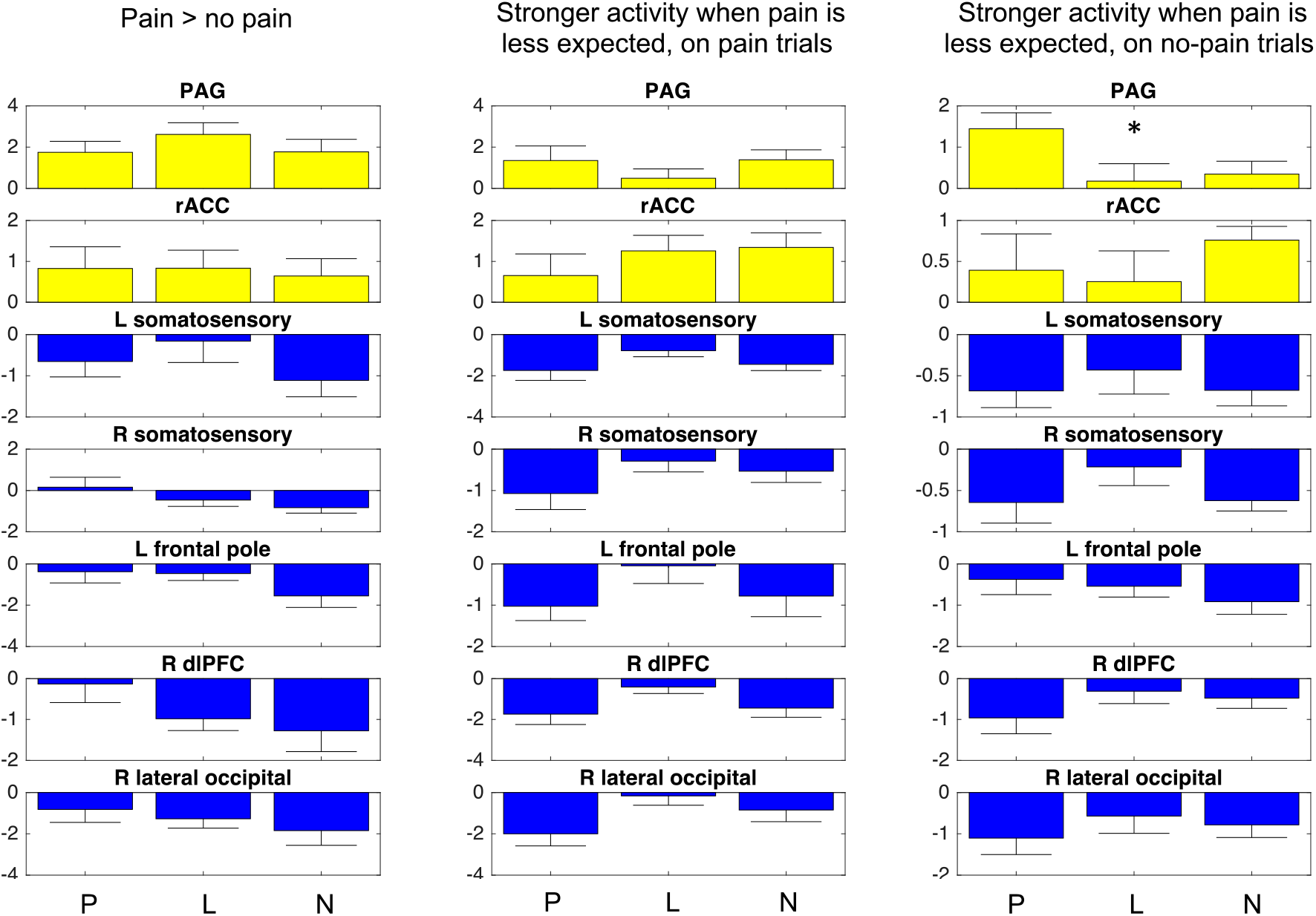
Activation for each of our three prediction-error contrasts, extracted from the clusters identified by our conjunction analysis, per treatment group. The three bars in each plot correspond to the placebo, levodopa and naltrexone group, respectively. There was only one significant treatment effect: The negative correlation with expected pain probability on no-pain trials (axiom 3) in the PAG cluster was stronger in the placebo group than in the drug groups (*p =* .04). As this result did not survive correction for multiple tests, and there were no significant treatment effects for the other prediction-error contrasts in any of the clusters, we conclude that our pharmacological manipulations did not affect prediction-error related brain activation. Error bars indicate standard errors. * p < .05 (uncorrected for multiple tests).

## Notes

### Competing Interest Statement

The authors have declared no competing interest.

